# The genetic architecture of recurrent segregation distortion in *Arabidopsis thaliana*

**DOI:** 10.1101/158527

**Authors:** Danelle K. Seymour, Eunyoung Chae, Burak I. Ariöz, Daniel Koenig, Detlef Weigel

## Abstract

The equal probability of transmission of alleles from either parent during sexual reproduction is a central tenet of genetics and evolutionary biology. Yet, there are many cases where this rule is violated. Such violations limit intraspecific gene flow and can facilitate the formation of genetic barriers, a first step in speciation. Biased transmission of alleles, or segregation distortion, can result from a number of biological processes including epistatic interactions between incompatible loci, gametic selection, and meiotic drive. Examples of these phenomena have been identified in many species, implying that they are universal, but comprehensive species-wide studies of segregation distortion are lacking. We have performed a species-wide screen for distorted allele frequencies in over 500 segregating populations of *Arabidopsis thaliana* using reduced-representation genome sequencing. Biased transmission of alleles was evident in up to a quarter of surveyed populations. Most populations exhibited distortion at only one genomic region, with some regions being repeatedly affected in multiple populations. Our results begin to elucidate the species-level architecture of biased transmission of genetic material in *A. thaliana*, and serve as a springboard for future studies into the basis of intraspecific genetic barriers.

## Introduction

At the genetic level, evolution is the change in the frequency of allelic variants over time. While in many cases the strength of selection is too low for these changes to be detected within a few generations, a unique opportunity to directly study such changes is offered in cases where selection coefficients are high. In such a situation, competition between alleles can be seen already in the distribution of heterozygous progeny (a/A). It is manifested as a deviation from the 1:2:1 Mendelian ratio of diploid genotypes (a/a, a/A, A/A), termed allelic or segregation distortion. Deviation from this ratio has important implications for population dynamics. Favoring inheritance of an allele from one grandparent over that from the other grandparent implies that certain genotypic combinations may be unfit (FISHMAN AND SAUNDERS 2008; PHADNIS AND ORR 2009; MCDERMOTT AND NOOR 2010). Depending on the underlying mechanism, complete obstruction to the free flow of genetic information can be an irreversible step on the path towards speciation (reviewed in (PRESGRAVES 2010)).

Segregation distortion, which is quite commonly observed in nature, can be the result of deleterious epistatic interactions between incompatible loci, gametic selection, or meiotic drive (reviewed in (LYTTLE 1991)). Perhaps epistatic interactions of the Bateson-Dobzhansky-Muller type are the best-studied examples (ORR 1996). Alone, the mutations causal for incompatibilities in progeny are innocuous in their native genetic environment. But when combined, their reduced fitness or lethality removes incompatible genotypic combinations from the population. Examples of two-locus incompatibilities have been identified in and between several eukaryotic species (reviewed in (ORR AND PRESGRAVES 2000; BOMBLIES AND WEIGEL 2007; RIESEBERG AND WILLIS 2007)) and causal loci are frequently associated with fast molecular evolution (reviewed in (ORR AND PRESGRAVES 2000; BOMBLIES AND WEIGEL 2007)). Likely, strong epistatic incompatibilities are a common topic in the literature not only due to their role in speciation, but also because they are easy to detect.

Far fewer examples of meiotic drive and gametic selection have been characterized. Meiotic drive refers to the preferential inheritance of one chromosome during meiosis and is most easily discovered during female gametogenesis (SANDLER *et al.* 1959), as only one of the four meiotic products will become the egg nucleus. This creates the opportunity for “selfish” loci to position themselves favorably so that they their transmission is favored in the next generation. Some known examples of female drive involve changes in either centromeric or other heterochromatic regions (MALIK AND HENIKOFF 2002; FISHMAN AND SAUNDERS 2008), possibly favoring transmission of the drive chromosome by increasing its affinity for the meiotic machinery (STURTEVANT AND DOBZHANSKY 1936; RHOADES 1942; SANDLER *et al.* 1959; HARTL *et al.* 1967; RHOADES *et al.* 1967; DUNN AND BENNETT 1968; ZIMMERING *et al.* 1970; FISHMAN AND SAUNDERS 2008). Many known drive loci are located on sex chromosomes (especially in various *Drosophila* species) and are associated with inversions or other cytological changes (STURTEVANT AND DOBZHANSKY 1936; ZIMMERING *et al.* 1970; FISHMAN AND SAUNDERS 2008). Drive loci on sex chromosomes are more readily identified because they alter the sex ratio, which is easily noticed without molecular biology assays.

Transmission biases arising after formation of the haploid gametes are classified as instances of gametic selection. Due to the differences of male and female gametogenesis, gametic selection can be more easily detected in males. Sperm is produced from all four meiotic products, and each of these haploid sperm cells can compete for the ability to fertilize the ovule. A classic example of gametic selection involves growth of the pollen tube that delivers the male gametes of plants (SNOW *et al.* 2000). For example, differential pollen tube growth can improve the reproductive success of the genotype that elongates more quickly.

A few instances of segregation distortion are well understood, but knowledge of the species-wide prevalence of the phenomenon is mostly missing. Despite the apparent ubiquity of segregation distortion, it is unclear how often epistatic incompatibilities, gametic selection, or meiotic drive are the cause. In *A. thaliana*, segregation distortion due to partially or fully recessively acting alleles has been observed repeatedly in different experimental population designs (LISTER AND DEAN 1993; MITCHELL-OLDS 1995; ALONSO-BLANCO *et al.* 1998; LOUDET *et al.* 2002; WERNER *et al.* 2005; SIMON *et al.* 2008; TÖRJÉK *et al.* 2008; BALASUBRAMANIAN *et al.* 2009; SALOMÉ *et al.* 2012). The largest published study to date in *A. thaliana* examined segregation distortion in 17 F_2_ populations, over half of which exhibited evidence of distortion (SALOMÉ *et al.* 2012). Although *A. thaliana* is typically a self-fertilizing species, outcrossing in nature can be quite common, implying that opportunities for unequal transmission shaping genetic diversity exist (BOMBLIES *et al.* 2010). On the other hand, the preference for inbreeding creates a system sensitized for detection of intraspecific distortion, since accessions collected from nature are typically homozygous throughout the genome. Cross-fertilization between accessions removes an allele from its native, homozygous context, thus creating an opportunity for biased transmission, which in turn makes *A. thaliana* an ideal system for the identification of preferentially inherited loci.

We have surveyed over 500 segregating F_2_ populations for segregation distortion in order to characterize the contribution of biased transmission to the generation of intraspecific genetic barriers. Segregating F_2_ populations were derived from intercrossing 80 distinct, resequenced *A. thaliana* accessions spanning the Eurasian range of the species (CAO *et al.* 2011). For this large survey, populations were genotyped in pools using reduced-representation high-throughput sequencing to estimate allelic ratios. In addition to documenting the prevalence of segregation distortion in *A. thaliana*, we have also begun to dissect the population-wide genetic architecture of segregation distortion. The crosses and genomic regions we have characterized provide a platform with which to dissect the relative contribution of deleterious epistatic interactions, male gametic selection, and female drive meiotic to biased inheritance.

## Materials and Methods

**Germplasm.** The F_2_ populations were generated by intercrossing 80 natural *Arabidopsis thaliana* accessions with whole-genome resequencing information (CAO *et al.* 2011). Intercrossing was facilitated by induced male sterility which was achieved by artificial miRNA (amiR) mediated knock-down of the floral homeotic gene APETALA3 (AP3) (CHAE *et al.* 2014). One half of F1 plants were transgene-free and able to produce F2 progeny through self-fertilization, as each original female grandparent was hemizygous for the amiR transgene. In total, 583 F_2_ populations were generated using 67 of the 80 natural accessions as the female grandparent. All 80 accessions were used as the male grandparent and, on average, each grandparent contributed to 14.7 F2 populations. Germplasm information can be found in Table 1 and grandparental seed availability is listed in Table S1.

**Growth conditions.** At least 300 individuals from each F_2_ population were sown onto 0.5x MS medium (0.7% agar; pH 5.6). Prior to plating, seeds were gas sterilized for 16 hours using 40 ml of household bleach (1-4%) and 1.5 ml of concentrated HCl. Seeds were stratified at 4°C in the dark for 8 days and then plates were shifted to 23°C long day conditions (16 h light:8 h dark). After 5 days, seedlings were harvested in bulk and flash frozen in liquid nitrogen.

**DNA extraction and GBS library preparation.** DNA was extracted from each pool of F_2_ individuals using a CTAB procedure (2% CTAB, 1.4 M NaCl, 100 mM Tris (pH 8), 20 mM EDTA (pH 8)) (SPRINGER 2010). DNA integrity was confirmed by gel electrophoresis, and DNA quantification was performed using the Qubit fluorimeter (Qubit BR assay) (Thermo Fisher Scientific, Waltham, MA). For library preparation, 300 ng of each DNA sample were diluted in 27 μl. Restriction enzyme-mediated reduced-representation libraries were generated using KpnI, which is predicted to cleave the *A. thaliana* reference genome into 8,366 fragments. The library preparation protocol is detailed in (ROWAN *et al.* 2017). Briefly, DNA was digested and then ligated to barcoded adapter sequences with sticky ends complementary to the KpnI cleavage site. After ligation, 96 barcoded samples were pooled and then sheared using the Covaris S220 instrument (Covaris, Woburn, MA). Next, end-repair, dA-tailing, a second universal adapter ligation, and PCR enrichment were performed using the Illumina compatible NEBNext DNA Library Prep Master Mix Set (NEB, Ipswich, MA). Library quality was determined using the Agilent 2100 Bioanalyzer (DNA 1000 kit) (Agilent, Santa Clara, CA) and libraries were normalized (10 nM) based on library quantification (ng/μl) and mean fragment length. Sequencing was performed on the Illumina HiSeq 2000 (Illumina, San Diego, CA). Adapter sequences can be found in (ROWAN *et al.* 2017).

**SNP identification and allele frequency estimation.** SHORE software (v0.9.0) (OSSOWSKI *et al.* 2008) was used for all analyses described in this section. Sequencing reads were barcode sorted and quality filtered. During quality filtering the restriction enzyme overhang was also trimmed using SHORE import. Reads for each bulked population were then aligned to the TAIR10 reference genome allowing for two mismatches using SHORE mapflowcell. After alignment, SNPs were called with SHORE qVar using default parameters. Read counts for both the reference and non-reference base were extracted for each polymorphic position. SNPs were filtered further using the grandparental whole-genome information and read counts for the female grandparental allele were output only for positions expected to be segregating between the two initial grandparents based on the resequencing data (CAO *et al.* 2011). The allele frequency of the female grandparental allele was calculated for each polymorphic position as the number of reads containing the female grandparental allele divided by the total number of reads covering that position.

**Modeling of allele frequency and significance testing for allelic distortion.** High read coverage was sought for each library to enable accurate allele frequency estimation. The realized median coverage of the population bulks was 78x. The distribution of read coverage per library is shown in Fig S1A.

Even with high read coverage, allele frequency estimates were still noisy. To generate accurate allele frequency estimates, the allele frequency was modeled in 5 Mb sliding windows (0.5 Mb steps). We used a beta-binomial model to account for variation in the true allele frequency as well as stochastic variation that arises from read sampling. From the optimized model we extracted the alpha and beta parameters from each genomic window. These parameters describe the shape of the probability distribution in each window, and from these parameters the mean allele frequency as well as the 95% confidence intervals were estimated. Using these estimates, a nonparametric statistical test was performed to assess whether the allele frequency estimates were significantly different from 50%, the expected frequency for non-distorted genomic regions. A false discovery correction (FDR) was performed to account for the number of genomic windows tested per population (n = 240). After allele frequency estimation, quality control measures culled low quality bulks. Populations were excluded from subsequent analysis for the following reasons: 1) having a genomewide average allele frequency greater than 0.75, 2) exhibiting either confidence intervals (CI) larger than 0.40 or noisy confidence intervals across the genome (standard deviation of CI width greater than 0.15), or 3) displaying three or more chromosomes with windows that did not attain model convergence. After quality control, 492 populations remained for subsequent analyses.

**Identification of distorted regions.** Two thresholds were used to identify significantly distorted genomic windows. The first approach utilized p-value estimates from the non-parametric statistical test performed on each window. False discovery rate (FDR) corrections were applied to account for the number of tested genomic windows (n = 240, p < 0.05). Distorted populations were required to have at least five adjacent genomic windows on the biased chromosome with significant FDR corrected p-values. Populations with statistically significant segregation distortion are listed in Table 1.

The second, less conservative approach identified outliers by calculating Z-scores for each genomic window relative to the mean allele frequency of all surveyed F2 populations (0.5029). Allele frequencies for each window were derived from the beta-binomial model predictions. Genomic windows with allele frequency estimates greater than 2.5 times the population-wide standard deviation (0.0382) were considered to be distorted. A distorted F_2_ population was required to contain five genomic windows with significant Z-scores on the chromosomes containing the locus of interest. Distorted populations identified using extreme Z-scores are listed in Table 1.

**Interval identification using whole-genome resequencing.** Six F_2_ populations displayed severe distortion at one of six distinct genomic regions (Fig S4). 1,500 individuals were sown from each of these six populations onto 0.5x MS medium (0.7% agar; pH 5.6) as described for the initial screen. DNA was extracted from each population bulk using a standard CTAB preparation (2% CTAB, 1.4 M NaCl, 100 mM Tris (pH 8), 20 mM EDTA (pH 8)). Illumina TruSeq libraries were prepared according to manufacturer’s guidelines using 1 μg of starting material per population. Libraries were sequenced on an Illumina HiSeq 3000 instrument (Illumina, San Diego, CA). Twenty-one nucleotide long k-mers were identified directly from the short reads using jellyfish (v2.2.3) (MARCAIS AND KINGSFORD 2011) with the following arguments: -m 21 -s 300M -t 10 -C. Not only does jellyfish identify all unique k-mers, but it also calculates the occurrence, or coverage, of each k-mer. The distribution of 21-mer coverages is shown in Figure S3 for each population. 21-mers found in only one of the two grandparental genomes (coverage < 25X) were aligned to the TAIR10 genome using bwa aln (LI AND DURBIN 2009. Only perfect matches were allowed. A 1 Mb sliding window (50 kb steps) was used to plot the 21-mer coverage across the distorted chromosome in each population. Regions of the genome with reduced coverage of 21-mers are located within the candidate interval (Fig 6B, S4). Interval boundaries were delineated by merging all windows with values within 1x coverage of the minimal window in the candidate region.

**Interval identification for distortion bulked segregant analysis.** Bulked segregant analysis (MICHELMORE *et al.* 1991) was used to narrow the candidate intervals for Star-8, ICE49, and ICE63. Sequencing reads from the original screen were combined for all distorted populations sharing the grandparent of interest, resulting in a distorted bulk. Those that shared the grandparent, but did not exhibit distortion, were combined separately, resulting in a normal bulk. Positions segregating between the grandparent of interest and all other members of the bulk were identified. The positions segregating in the distorted bulk are not shared with those segregating in the normal bulk. By combining reads from multiple populations, a median of 806 to 1135x coverage was achieved at each segregating position. Candidate intervals were calculated from the maximally distorted position to any flanking segregating site that was within 5% of the peak allele frequency (Table 2).

**Material and data availability:** Seeds for grandparental lines are available from the Arabidopsis Biological Resource Center (ABRC) or the European Arabidopsis Stock Center (NASC); stock identifiers are listed in Table S1. The source code to generate allele frequency estimates and the raw allele frequencies for each F2 population are located in the following github repository: https://github.com/dkseym/F2_Segregation_Distortion.

## Results

**Frequent segregation distortion in intraspecific *A. thaliana* F_2_ populations.** The incidence of segregation distortion, a molecular signature of genetic conflict, was surveyed in 583 F2 populations generated from naturally inbred accessions that represent much of the Eurasian genetic diversity in *A. thaliana* (CAO *et al.* 2011). The studied F2 populations were derived from crosses between 67 accessions used as female and male grandparents, and a further 13 that were used only as male grandparents (CAO *et al.* 2011). The number of crosses performed per accession ranged from 3 to 34, with a median of 14 F_2_ populations generated from each grandparent.

F2 seeds were sown on plates, stratified at 4°C to break dormancy, and then grown for five days in 23°C long days. At least 300 individuals per F_2_ population were harvested in bulk for genome-wide genotyping-by-sequencing (GBS), implemented as restriction enzyme-mediated reduced-representation sequencing. Based on previous reduced-representation approaches (BAIRD *et al.* 2008; MONSON-MILLER *et al.* 2012), a custom protocol was developed to adapt this method to the specific requirements of our system. Accurate allele frequency estimate in bulks requires high sequencing coverage at each segregating site. The selected restriction enzyme, KpnI, cuts infrequently in the *A. thaliana* genome, allowing high coverage to be achieved for a portion of the genome, about 1%, with moderate sequencing effort. Whole-genome SNP information was available for the inbred grandparents (CAO *et al.* 2011), facilitating identification of informative sites. We attained an average of 78x coverage per F2 population (Fig S1A), and an average of 2,509 sites were segregating in any given population (Fig S1B).

Regions displaying significant segregation distortion, as indicated by deviation from the expected 1:1 ratio of grandparental alleles, were identified by modeling the allele frequency in 5 Mb sliding windows, with 0.5 Mb steps. Using the beta-binomial model estimates of allele frequencies together with the confidence intervals of the estimates, a non-parametric statistical test was performed in each window. In total, 62 populations exhibited regions of significant segregation distortion after false discovery rate (FDR) correction for the number of tested windows (n = 240, p < 0.05). When considering only the 492 populations passing quality control measures, 62 (12.6%) of these were found to harbor genomic regions with significant distortion (Fig S2). This is a rather conservative estimate of the incidence of segregation distortion in our crosses, because the ability to detect significant distortion is highly dependent on the size of the confidence interval estimates (i.e., the coverage of each population).

To generate a less conservative estimate of the number of distorted regions, we also used a Z-score outlier approach. Any region with allele frequencies greater than 2.5 standard deviations from the combined population mean was considered to be distorted. This less conservative approach identified 122 (24.8%) of the 492 populations with at least a single distorted region (Fig 1). All regions identified via the FDR method were also detected using the Z-score outlier approach.

**Figure 1.**
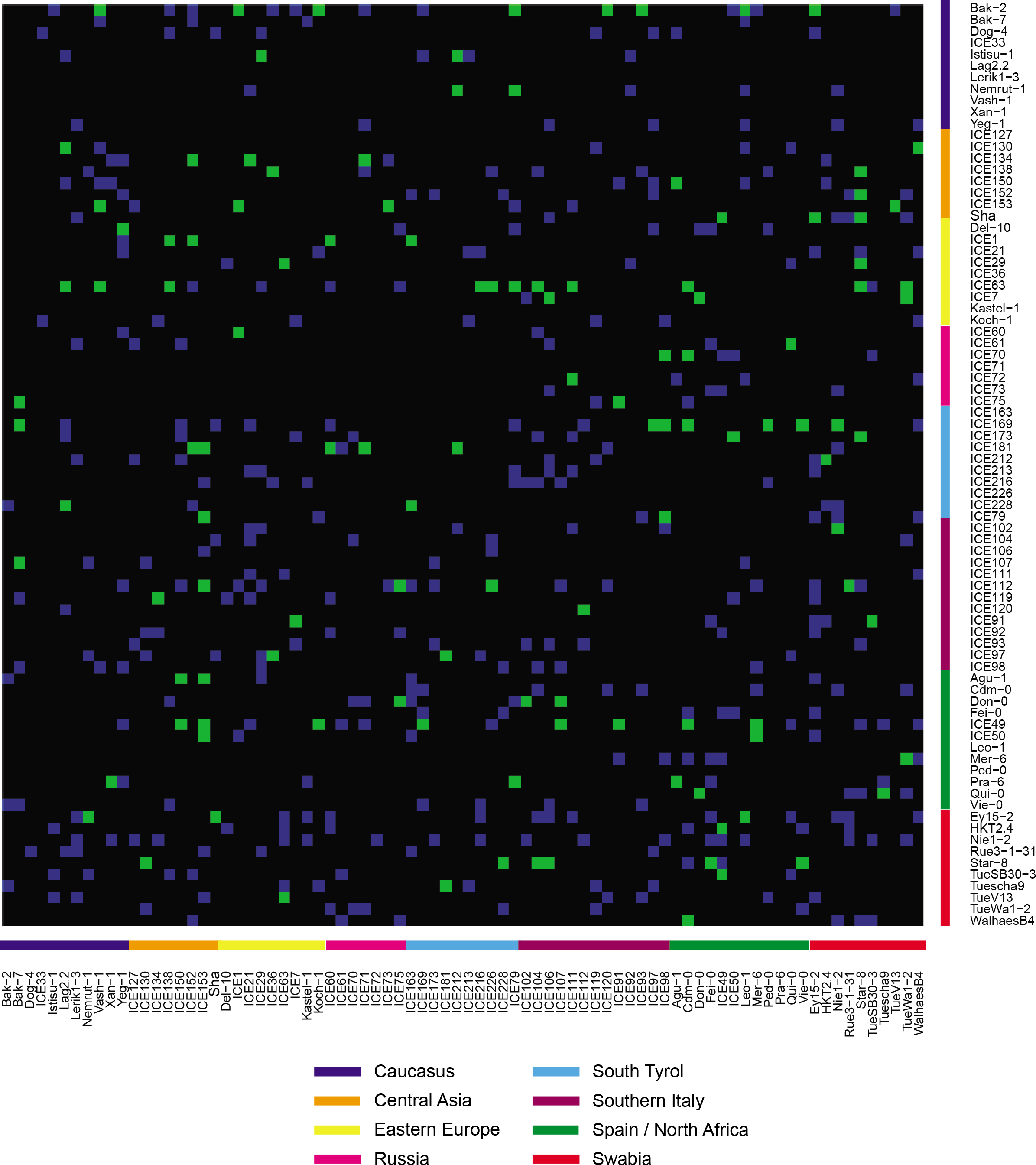
**Z-score estimated segregation distortion is evident in a wide range of crosses.** Genotypic combinations surveyed in this F_2_ screen are shown in blue, and populations with significant segregation distortion based on Z-score metrics in green. Grandparental accessions are ordered by the geographic location of their collection (CAO *et al.* 2011). Female grandparents are located on the y-axis and male grandparents on the x-axis.

An example of a chromosome with a distorted region that was identified using both methods is shown in Figure 2. Although we did not screen the complete diallel of possible F2 combinations, we did survey populations that sampled a large fraction of the genetic space covered by the 80 founders (Fig 1, Fig S2). That segregation distortion is evident in up to 24% of surveyed F2 populations suggests that intraspecific genetic barriers are much more common than previously anticipated.

**Figure 2.**
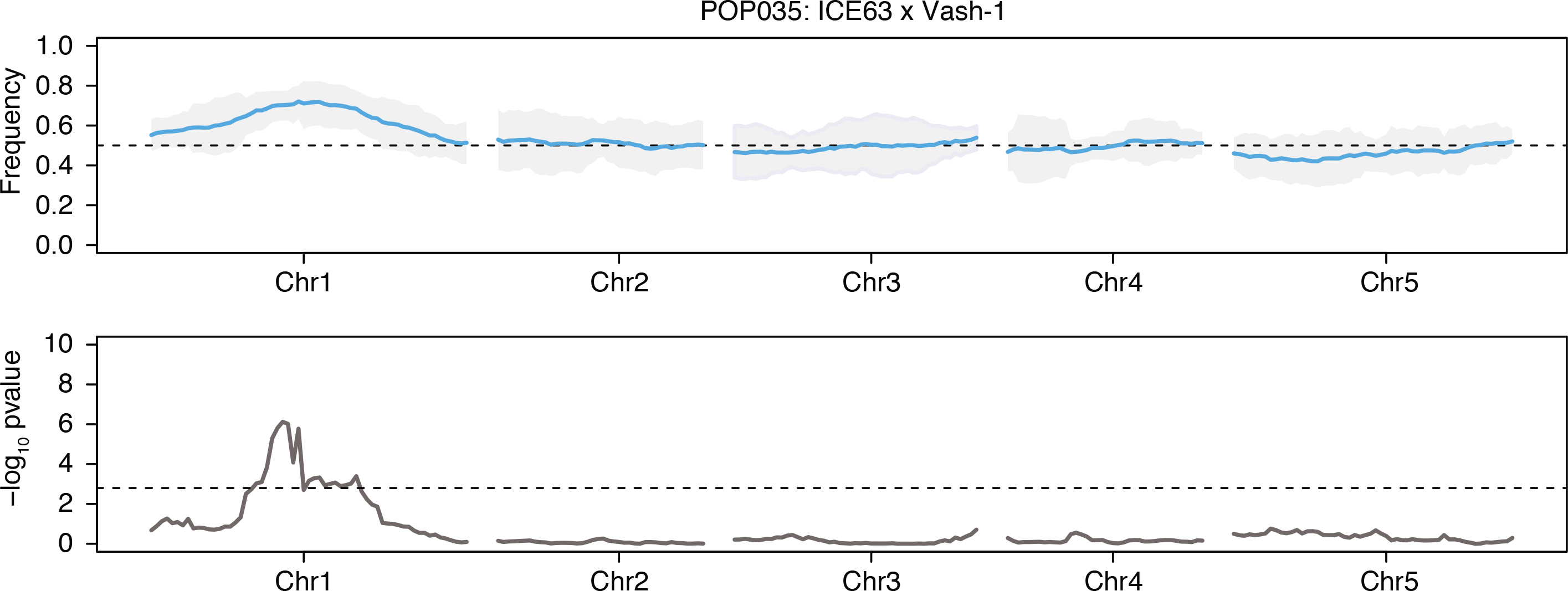
**A representative F_2_ population, POP035 (ICE63 x Vash-1), with significant segregation distortion.** Distortion in this population was detected with both thresholds (FDR and Z-score outlier). (A) The beta-binomial modeled allele frequency (blue) across each chromosome is plotted in the upper panel. 95% confidence intervals are indicated by the shaded grey area and the expected frequency of 0.5 is marked by the dashed black line. (B) The -log™ of the p-value derived from the non-parametric statistical test. The dashed black line in this panel represents the FDR corrected (n = 240) significance threshold (p < 0.05).

**The dynamics of segregation distortion in *A. thaliana.*** The genetics of segregation distortion is dictated by the biological process driving the observed non-Mendelian inheritance. To understand the relative contribution of different processes such as genetic incompatibility, meiotic drive, and gametic selection, we determined how many genomic regions showed segregation distortion in our data set.

Regardless of identification method – FDR or Z-score outlier –, the majority of populations exhibited distortion at only a single locus (Fig 3A). If classical Bateson-Dobzhansky-Muller genetic incompatibilities were driving segregation distortion in our populations, we would expect two distorted regions per population, unless the responsible loci were linked. We also found that distortion occurs on all five chromosomes, although distorted regions are most frequently located on chromosome 1 (Fig 3B).

**Figure 3.**
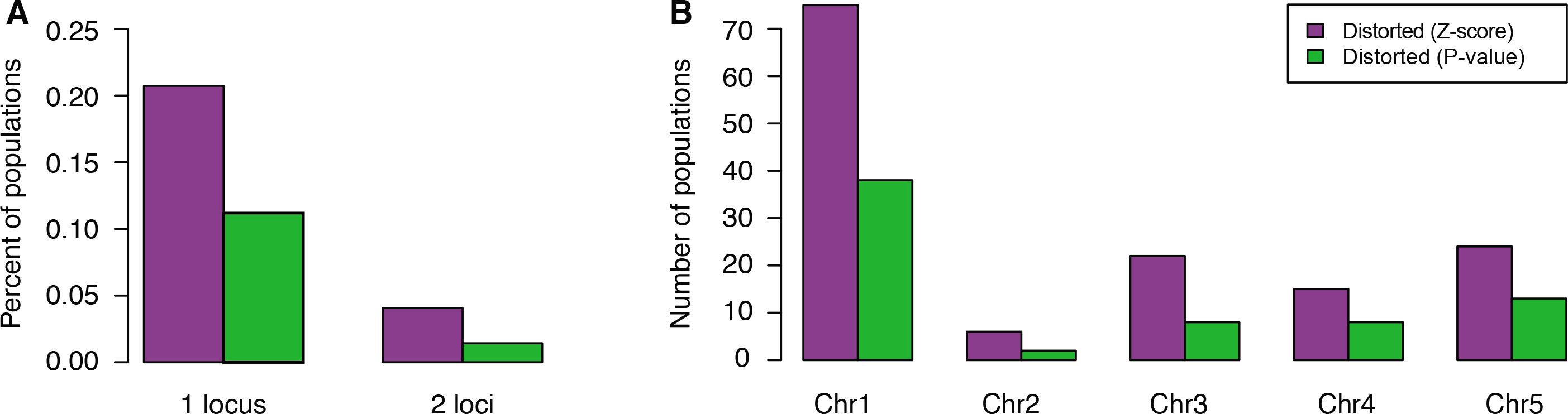
**Genomic properties of distorted loci.** (A) The fraction of surveyed F_2_ populations that exhibited segregation distortion at either one or two genomic loci. (B)The number of populations containing distorted loci that reside on each of the five *A. thaliana* chromosomes.

The alleles in distorted regions that are favored to be inherited are derived from many grandparental accessions. Of the 80 accessions used as founders, over 50 gave rise to F_2_ populations exhibiting significant segregation distortion. Some grandparents were especially notable, such as Star-8. Regions with alleles contributed by Star-8 were distorted in 60% of F_2_ populations (40% for the FDR threshold) (Fig 4A,B).

**Figure 4.**
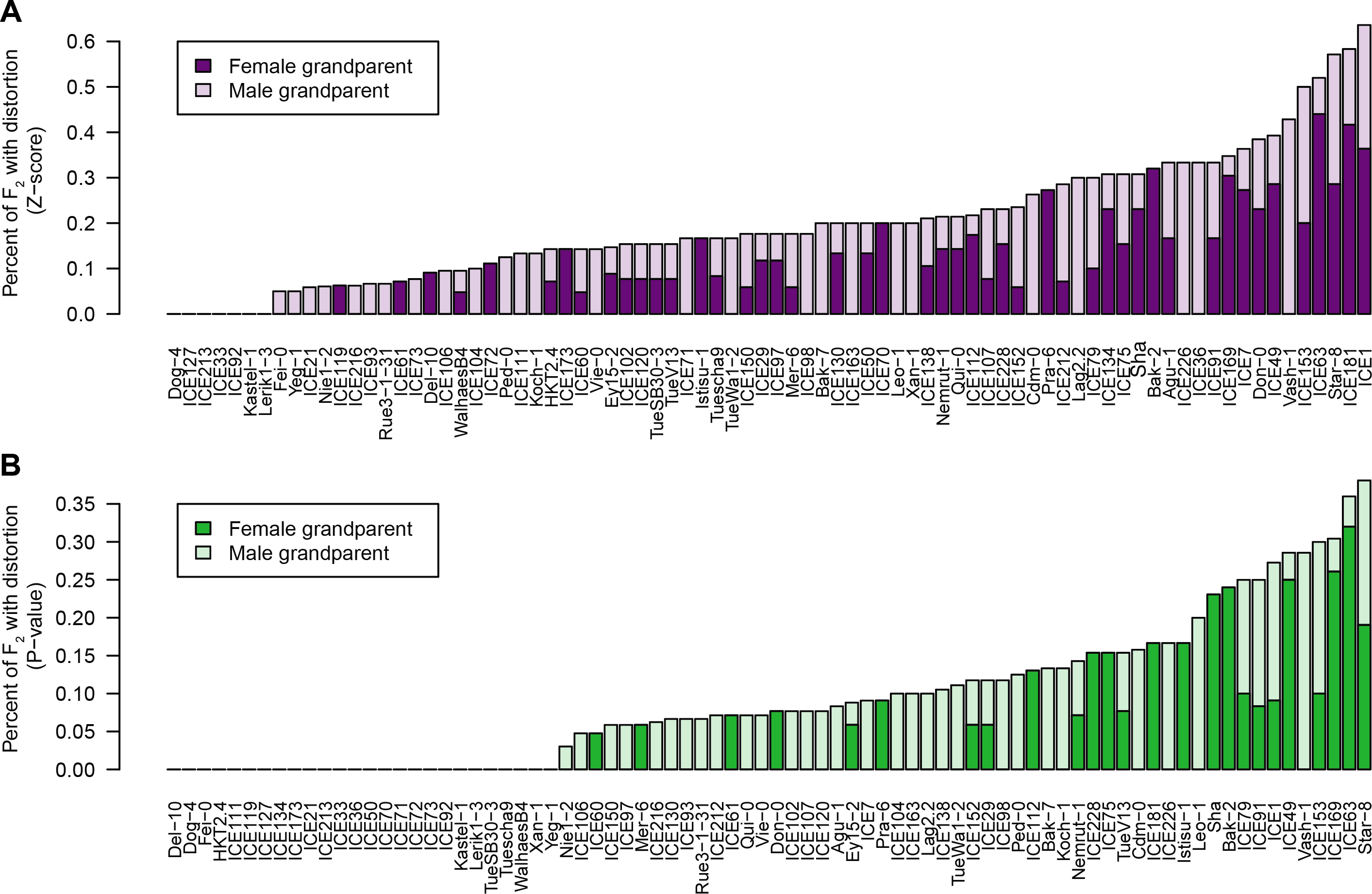
**Many grandparental accessions contributed biased alleles.** Each grandparent contributed its genetic material to a median of 14 distinct F_2_ populations. Plotted is the fraction of F2 populations with one shared grandparent that are significantly distorted as measured either by (A) because of FDR corrected deviation from beta-binomial modeled allele frequencies, or (B) 2.5x Z-score deviation.

If genetic barriers are primarily caused by genetic drift as individuals diverge from a common ancestor, we would expect more distantly related accessions to give rise to distortion more frequently (LEPPALA *et al.* 2013). As the grandparental accessions had been sampled from eight geographic regions representative of Eurasian genetic diversity (CAO *et al.* 2011), we were able to test if genetic diversity between the two F_2_ grandparents was correlated with the probability of segregation distortion. We found no significant difference between the genetic distances of grandparents of distorted populations compared to grandparents of non-distorted populations (Wilcoxon rank-sum test, 1% significance threshold; p=0.03 [Z-score outlier distortion list], p = 0.11 [FDR list]) (Fig 5A,B). That genetic diversity is not a strong predictor of segregation distortion suggests that genetic drift, which becomes more notable after longer periods of separation, is not necessarily the most important driver of intraspecific genetic barriers in *A. thaliana.*

**Figure 5.**
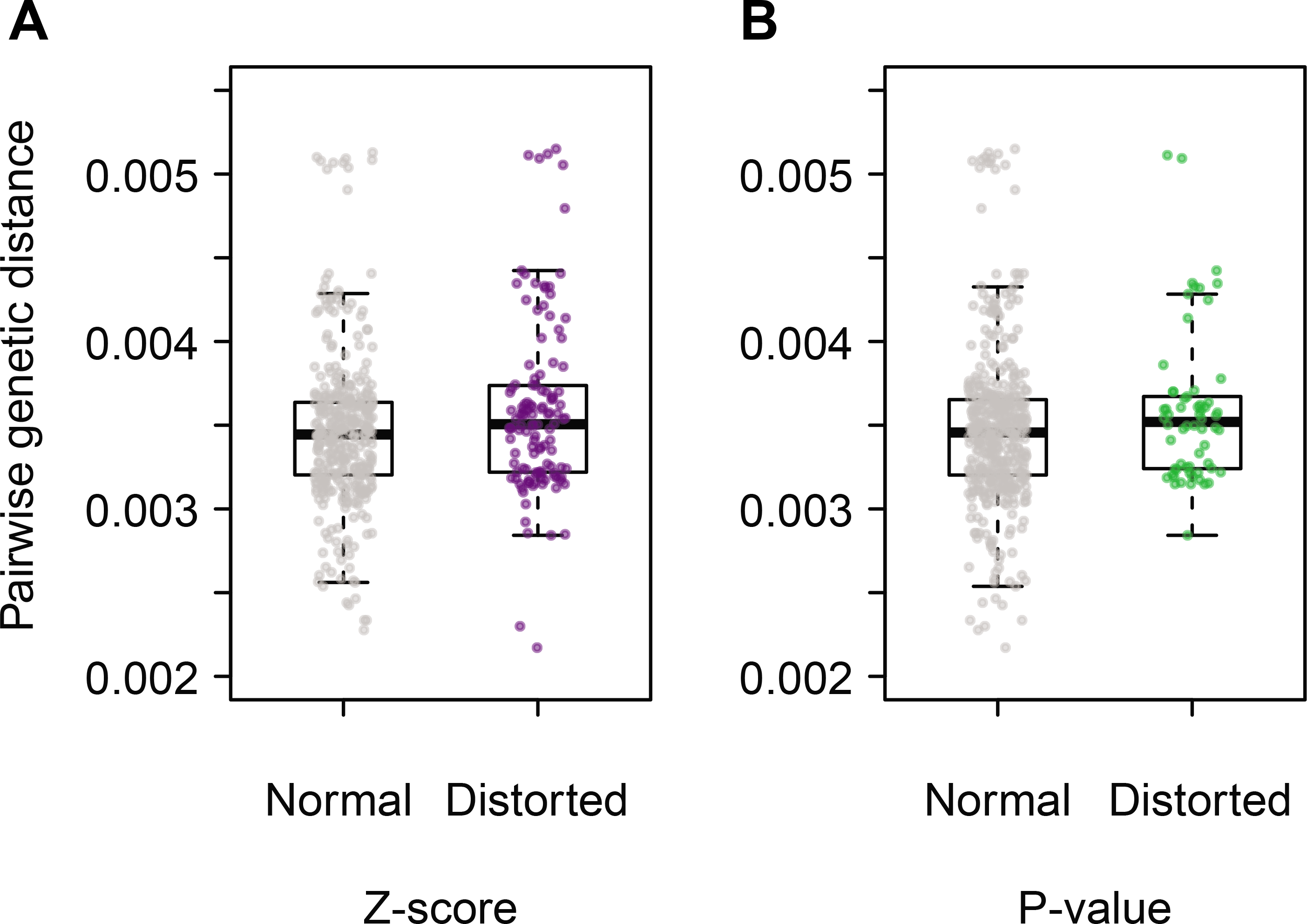
**Genetic distance between grandparental accessions is not predictive of biased allelic transmission.** A box plot of genetic distances between the grandparental accessions of normal (grey) and distorted (colored) F2 populations. At a significance threshold of p < 0.01, the genetic distances between grandparents of distorted populations determined from FDR corrected deviation (A) or 2.5x Z-score deviation (B) is not significantly different from that of normal populations (Wilcoxon rank sum test). Genetic distance was calculated as the number of segregating sites over the number of interrogated sites. All positions were required to have complete coverage across all 80 grandparental accessions.

**Refining candidate intervals surrounding distorted loci.** To begin to understand which processes are responsible for the observed segregation distortion, we sought to define the minimal size of distorted genomic intervals. Genotyping F2 individuals in bulk enabled screening of a large number of test populations, but without genotype information from individual segregants to estimate recombination breakpoints, most candidate regions are not much smaller than entire chromosome arms.

Since we did not know a priori which populations would be the most informative to study in detail, we designed two strategies to narrow the candidate regions to facilitate subsequent fine-mapping. First, we increased the density of informative markers about 200 fold by whole-genome resequencing of six populations with severe segregation distortion. We also increased the number of recombination events in these populations by analysis of 1,500 F2 individuals from each of the six populations. We sequenced these bulks to approximately 40x coverage. Although this coverage was lower than the average 78x coverage we had used in our GBS analyses, by integrating over multiple markers, together with the larger number of F_2_ individuals and thus recombination events, we expected this to substantially improve our power to delineate distorted regions.

Unfortunately, exploratory analyses indicated that the lower coverage at individual markers is accompanied by increased stochasticity in allele frequency estimates. We therefore took advantage of local linkage disequilibrium to diminish that noise. Short stretches of unique 21 nucleotide (nt) sequences (known as k-mers or 21-mers) were identified in the raw sequencing reads of each F_2_ population. Any 21-mer sequence shared between grandparents should occur at the average genome-wide coverage, and when we plotted 21-mer frequencies, we found a major found peak of 21-mer coverage around 40x, the average per-population whole-genome coverage, in all six populations, as expected (Fig 6A, S3). In contrast, 21-mers present in only one of the two parents should have approximately half as much coverage, and a second peak, resulting from a much smaller number of 21-mers, was apparent in all populations as well (Fig 6A, S3).

**Figure 6.**
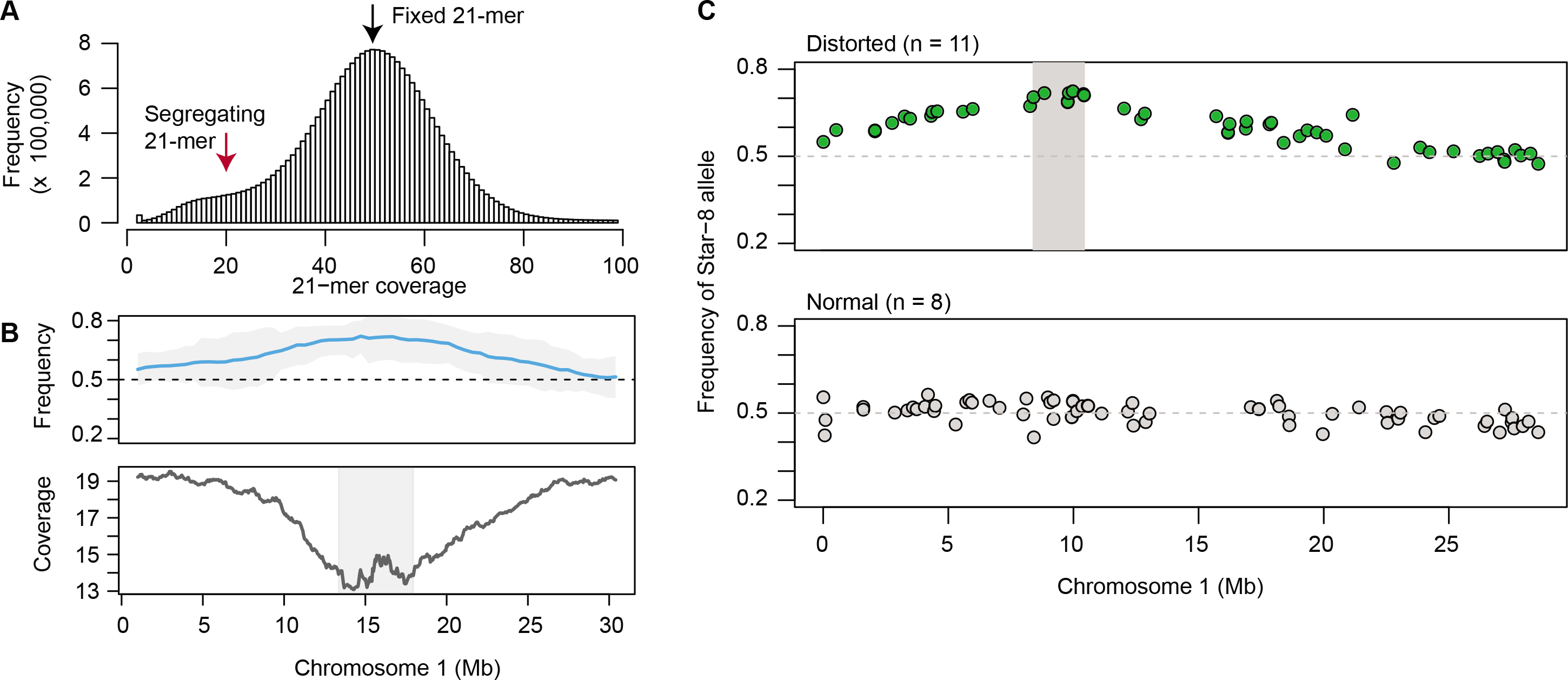
**Mapping intervals refined using k-mer coverage and bulked segregant analysis.** (A) The coverage of unique 21 nt k-mers is plotted for POP035 (ICE63 x Vash-1) after whole-genome resequencing. The first peak in coverage represents 21-mers found in only one of the two grandparents (red arrow), while the second, larger peak represents those sequences found in both (black arrow). (B) The upper panel displays the beta-binomial modeled allele frequency estimates (blue) and their 95% confidence intervals (grey) for POP035 as described in the legend for Figure 2. In the lower panel, the coverage of 21-mers unique to only one of the two grandparents (coverage < 25x) is plotted in 1 Mb sliding windows (50 kb steps). Coverage decreases in the candidate regions. Intervals (grey box) are defined by merging windows with values within 1x coverage of the minimal window in each population. (C) Bulked segregant analysis was performed for Star-8, an accession that repeatedly contributed distorted loci. Sequencing reads were combined for populations exhibiting distortion when crossed with Star-8, and for populations not exhibiting distortion when crossed to Star-8 (normal pool). A candidate interval (grey box) was obtained by merging all segregating positions within 5% of the maximal allele frequency.

To narrow down candidate intervals, we extracted 21-mers that were predicted to be present in only one of the two grandparents. Regions of the genome that are distorted should display a decrease in coverage of such grandparent-specific 21-mers near the causal locus. We used a sliding window approach (1 Mb windows, 50 kb steps) to calculate the average coverage of such 21-mers. Using this strategy, we were able to narrow the intervals surrounding four of the six candidate loci to less than 5 Mb, and in one case to 1.5 Mb (Table 2, Fig 6B, S4).

In a complementary approach, we sought to refine candidate regions by obtaining a more precise estimate of local allele frequency. To this end, we greatly increased sequencing coverage by combining information from cases with shared grandparents and the same distorted regions. As mentioned earlier, some grandparental accessions contributed alleles that were favored in multiple F2 populations. Star-8, ICE63, and ICE49 contributed alleles that were favored in at least 40% of crosses of these to other accessions (based on the Z-score outlier method), with the same regions being favored in all distorted populations sharing a particular grandparent. Using a bulked segregant analysis approach (MICHELMORE *et al.* 1991), we generated two pools of reads for each grandparent. One comprised the sequencing reads from all distorted populations and the other contained the sequencing reads from all non-distorted populations. The allele frequency of SNPs was calculated for sites segregating between the focal grandparent and all other accessions in either the distorted pool or the non-distorted pool.

A median coverage of at least 806x was achieved at each segregating site, vastly improving the accuracy of our estimates. For one grandparent, Star-8, we narrowed the interval to 2.0 Mb, in the middle of the top arm of chromosome 1, where recombination is high (Table 2, Fig 6C). This strategy was less successful for the other two grandparents, ICE63 and ICE49, likely because of the distortion being less strong in these cases as well as the location of the distorted regions near the centromere or on the distal chromosome arm, both parts of the chromosome where recombination is reduced (Table 2, Fig S5).

## Discussion

Despite the ubiquity of non-Mendelian segregation of alleles in natural populations, the genetic and molecular characterization of the responsible loci has been lagging (reviewed in (ZIMMERING *et al.* 1970; LYTTLE 1991; LYON 2003; FISHMAN AND SAUNDERS 2008; PHADNIS AND ORR 2009; HAMMOND *et al.* 2012; LARRACUENTE AND PRESGRAVES 2012). Such systems are most easily studied, when distortion is severe and differences in phenotypically distinct progeny classes are obvious (reviewed in (ZIMMERING *et al.* 1970)). Because sexual dimorphism is common, many of the earliest known cases were discovered because sex-ratio deviated greatly from 1:1 (reviewed in (ZIMMERING *et al.* 1970)). The effects of an allele that is preferentially inherited can be neutralized in a population by fixation of the allele or by the evolution of secondary modifiers. Many cases of segregation distortion were discovered in interspecific crosses (CAMERON AND MOAV 1957; MAGUIRE 1963; SIRACUSA *et al.* 1991; TAO *et al.* 2001; FISHMAN AND SAUNDERS 2008; ZANDERS *et al.* 2014), not because the phenomenon is more common in interspecific hybrids, but because the severity of distortion is extreme in the absence of species-specific modifiers, sometimes reaching fixation in only a generation or two (FISHMAN AND SAUNDERS 2008). The same loci responsible for segregation distortion in interspecific crosses may also underlie unexpected intraspecific segregation patterns. However, in intraspecific crosses, allele frequencies are often only perturbed by a few percent (LYTTLE 1991; FISHMAN AND SAUNDERS 2008), and without molecular genotyping techniques, such subtle allelic distortion will go mostly undetected.

Exploiting advances in sequencing and genotyping technology, we have been able to characterize segregation distortion in hundreds of intraspecific crosses. The identification of distorted regions greatly depends on sequencing coverage; in our system, a 10% deviation in absolute allele frequency becomes significant with approximately 100x sequence coverage, and more subtly distorted regions could be detected with even higher coverage. Similar pooled genotyping approaches have been used to identify distorted loci in other systems (CUI *et al.* 2015; BELANGER *et al.* 2016a; BELANGER *et al.* 2016b; WEI *et al.* 2017), illustrating the general power of this approach.

Although *A. thaliana* is self-compatible, outcrossing is reasonably common, and descendants of recent outcrossing events are easily found in wild stands of this species (BOMBLIES *et al.* 2010). By surveying a broad collection of germplasm for non-Mendelian inheritance, we could confirm that allelic distortion is a common feature of F_2_ populations, implying that allelic distortion has a major impact on shaping local genetic diversity. Not only do distorted loci segregate in up to a quarter of all F2 populations, but multiple genomic regions contribute to this phenomenon, with the degree of distortion varying both by population and by locus. Intraspecific distortion loci that have been identified in other systems typically occur at low population frequencies (HICKEY AND CRAIG 1966; PERKINS AND BARRY 1977; HIRAIZUMI AND THOMAS 1984; HAMMER *et al.* 1989; MCMULLEN *et al.* 2009; HOU *et al.* 2015; FRAGOSO *et al.* 2017), although there are exceptions, such as the tightly linked *zeel-1* and *peel-1* genes in *C. elegans* (SEIDEL *et al.* 2008; BEN-DAVID *et al.* 2017). The low frequency of the causal alleles has been hypothesized to result from antagonistic modifier loci having evolved in response to the fitness costs that are often linked to distortion loci (reviewed in (ZIMMERING *et al.* 1970; LYTTLE 1991)). In an interspecific *Drosophila* cross, the causal locus itself is responsible for both the distortion phenotype and for reduced gamete success (PHADNIS AND ORR 2009). We have found multiple cases of genomic regions that are distorted in one or very few population(s), suggesting that frequency of distortion alleles is often low in *A. thaliana* as well. This could be because these alleles are older, giving sufficient time for modifiers to evolve and rise to high frequency. If these are linked, we would not have detected them as separate genomic loci, as our mapping resolution was mostly chromosome arm scale.

Of particular interest are regions that are repeatedly distorted across many populations at extreme frequencies. For example, the Star-8 region on chromosome 1 is significantly favored in ∼50% of crosses, with this region being inherited by up to 70 or even 80% of the progeny. This could be an example of a young allele for which suppressors have not yet evolved, or it could be that the balance between fitness costs (if any) and the degree of distortion is stable at this frequency. The *D* locus in *Mimulus guttatus* is perhaps the best example of a stable distortion polymorphism, in this case caused by meiotic drive (FISHMAN AND SAUNDERS 2008). The measured degree of distortion at this locus (58:42) is predicted by the associated decrease in pollen viability (FISHMAN AND SAUNDERS 2008). This allele is segregating in about half of all individuals from a natural population (FISHMAN AND SAUNDERS 2008). Other instances of distortion loci segregating at intermediate frequencies are known, but the evolutionary dynamics of these cases are not as well characterized (reviewed in (ZIMMERING *et al.* 1970; LYTTLE 1991))

A peculiarity of allelic distortion in our panel of *A. thaliana* crosses is that in most cases, only a single genomic region is inherited in a non-Mendelian fashion. Classic meiotic drive systems consist of a distorter locus and a responder locus, with the two being almost always linked through an inversion or genetic rearrangement that reduces recombination between them (STALKER 1961; WU AND BECKENBACH 1983; SILVER 1985; LYTTLE 1991). As a result, classic drive loci are inherited as a single distorted genomic region. Our results are reminiscent of such cases, suggesting that several such loci are segregating in *A. thaliana*, although we cannot currently infer the number of genes in the mapping intervals responsible for segregation distortion.

Apart from meiotic drive, more conventional two-locus deleterious interactions conforming to the Bateson-Dobzhansky-Muller model of genetic incompatibilities can also perturb expected allelic (and genotypic) segregation ratios. A survey in *D. melanogaster* showed intraspecific genetic incompatibilities due to epistatic interaction between two (often unlinked) loci are not uncommon, with natural strains carrying an average of 1.15 incompatible loci (CORBETT-DETIG *et al.* 2013). Hybrid incompatibility is a common feature in both plants in animals, with many known cases of deleterious epistatic interactions between two nuclear loci segregating in *A. thaliana* (BOMBLIES *et al.* 2007; ALCÁZAR *et al.* 2009; BIKARD *et al.* 2009; VLAD *et al.* 2010; DURAND *et al.* 2012; CHAE *et al.* 2014; AGORIO *et al.* 2017; PLÖTNER *et al.* 2017). In our set of crosses, simultaneous distortion at two independent genomic regions was the exception. In our design, incompatible interactions would only be detectable if the F_1_ was fertile and dominance relationship between alleles was such that over 10% of the progeny did not give rise to seedlings. In other words, if both genes acted completely recessively and the doubly homozygous progeny failed to grow, they still would not be noticed in our segregation distortion scans. We note that even in cases where two independent genomic regions are significantly distorted in a single population, the absence of genotype data for individuals does not allow us to explicitly examine if these regions genetically interact. Although the nature of our experimental design has not yet revealed the species-wide architecture of partially or fully recessive epistatic interactions segregating in *A. thaliana*, this can be addressed in future studies by genotyping individuals instead of pools.

While a handful of classical segregation distortion loci has been molecularly characterized in detail (reviewed in (ZIMMERING *et al.* 1970; LYTTLE 1991; LYON 2003; LARRACUENTE AND PRESGRAVES 2012)), the molecular nature of most loci is still unknown. As a result, there is still much to be learned about the biological processes and evolutionary forces leading to uneven segregation, including whether such alleles are more likely to be evolutionarily old or young. For example, numerous cases of hybrid incompatibilities in *A. thaliana* are due to interactions between disease resistance genes, which have very divergent alleles, both because of rapid evolution and long-term balancing selection (BOMBLIES *et al.* 2007; ALCÁZAR *et al.* 2009; DURAND *et al.* 2012; CHAE *et al.* 2014). The fast evolution of centromeres and other satellite sequence repeats, a result of intragenomic conflict, has also been shown to cause or to be closely linked to allelic distortion (WU *et al.* 1988; FISHMAN AND SAUNDERS 2008; CHMATAL *et al.* 2014; MAHESHWARI *et al.* 2015). In our crosses, distorted regions often localized near centromeres.

Whether the conflict arises in interspecific or intraspecific crosses, it appears that natural selection, not genetic drift, is often responsible for the evolution of non-Mendelian inheritance. In support of this, we found little correlation between the degree of genetic differentiation between the grandparental accessions and the probability of observing allelic distortion in their progeny, in line with what has been seen in a much smaller panel of F2 populations (SALOMÉ *et al.* 2012).

To conclude, by surveying a large number of F2 populations descending from 80 genetically diverse grandparents, we were able to identify numerous genomic regions in *A. thaliana* that are not transmitted in a Mendelian fashion. Considering that our statistical power would not have allowed us to discover complete absence of genotypes resulting from higher-order epistatic interactions, it is likely that the regions we identified are only the tip of the iceberg. Notably, the majority of accessions tested contributed such distorted alleles, emphasizing the ubiquity of alleles that are unevenly transmitted. Together, these findings confirm the findings from other systems that genetic barriers segregating within wild species are more common that previously thought (SEIDEL *et al.* 2008; CORBETT-DETIG *et al.* 2013; HOU *et al.* 2015).

## Author contributions

D.K.S., D.K., E.C. and D.W. conceived the project. D.K.S., E.C. and B.I.A. generated the material and data. D.K.S. and D.K. analyzed the data. D.K.S. and D.W. wrote the manuscript with contributions from all authors.

## Acknowledgments

This work was supported by ERC AdG IMMUNEMESIS and the Max Planck Society.

## Tables

**Table 1. Germplasm information for surveyed F_2_ populations.** All crosses are listed, with those passing quality control (QC) indicated with a “1”. Similarly, “1” and “0” indicates whether distortion was detected using FDR significance testing of beta-binomial modeling of allele frequencies or Z-score deviation.

**Table 2. Candidate intervals for distorted loci.** ND, not determined.

## Supplemental tables

**Table S1. Germplasm identifiers.**

## Supplemental 791 figure legends

**Figure S1.**
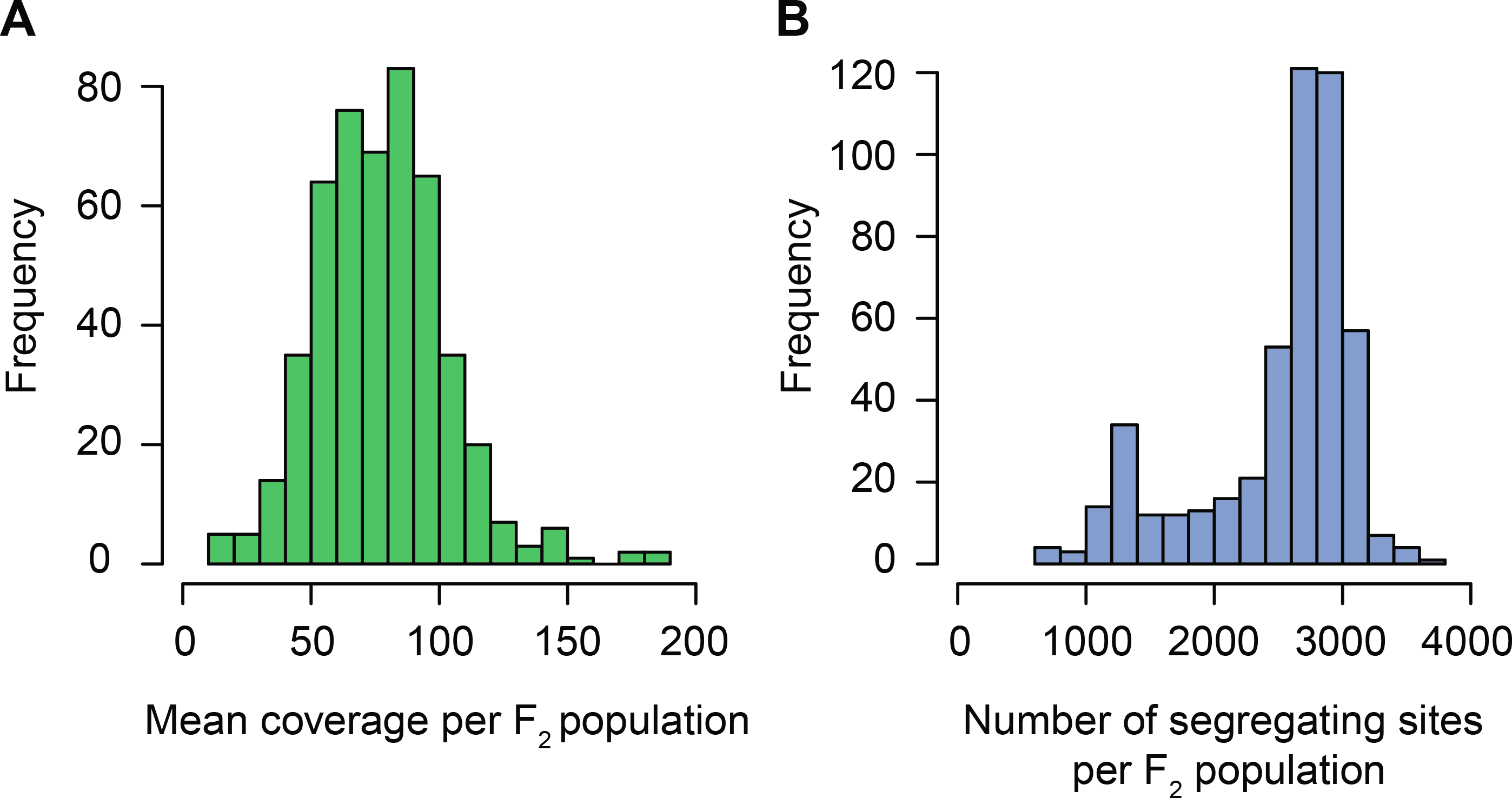
**Reduced-representation sequencing reliably enriches for 1% of the** *A. thaliana* **genome.** (A) Mean sequencing coverage at sites segregating in each F_2_ population. (B) Number of sites segregating in each F2 population. The mean observed number of segregating sites (2,500) is comparable to the expected number of segregating sites derived from previously published resequencing data (CAO *et al.* 2011).

**Figure S2.**
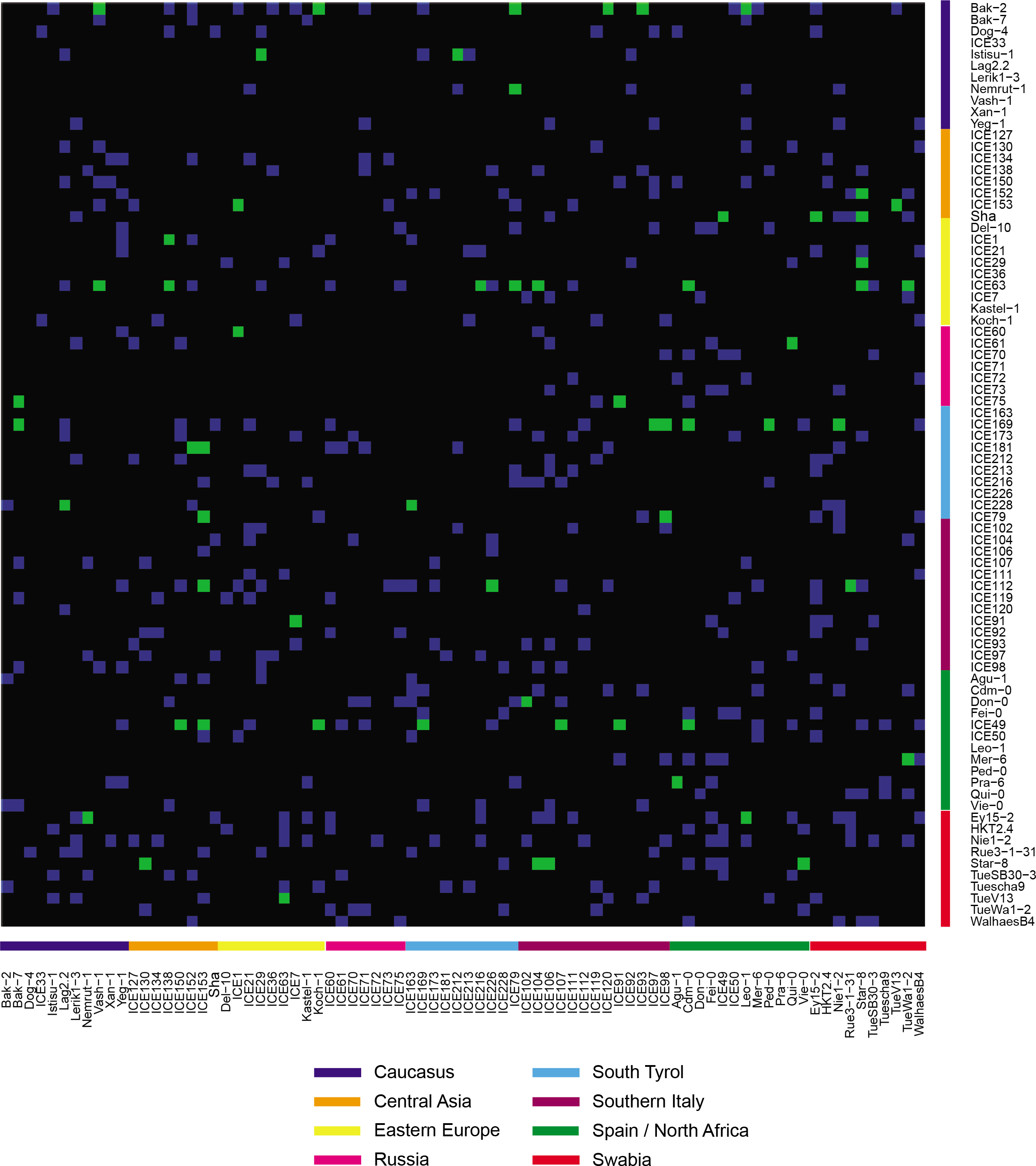
**Statistically significant segregation distortion is evident in a wide range of crosses.** Genotypic combinations surveyed in this F2 screen are shown in blue, and populations with significant segregation distortion based on non-parametric statistical tests of beta-binomial modeled allele frequencies in green. Grandparental accessions are ordered by the geographic region of their collection (CAO *et al.* 2011). Female grandparents are located on the y-axis and male grandparents on the x-axis.

**Figure S3.**
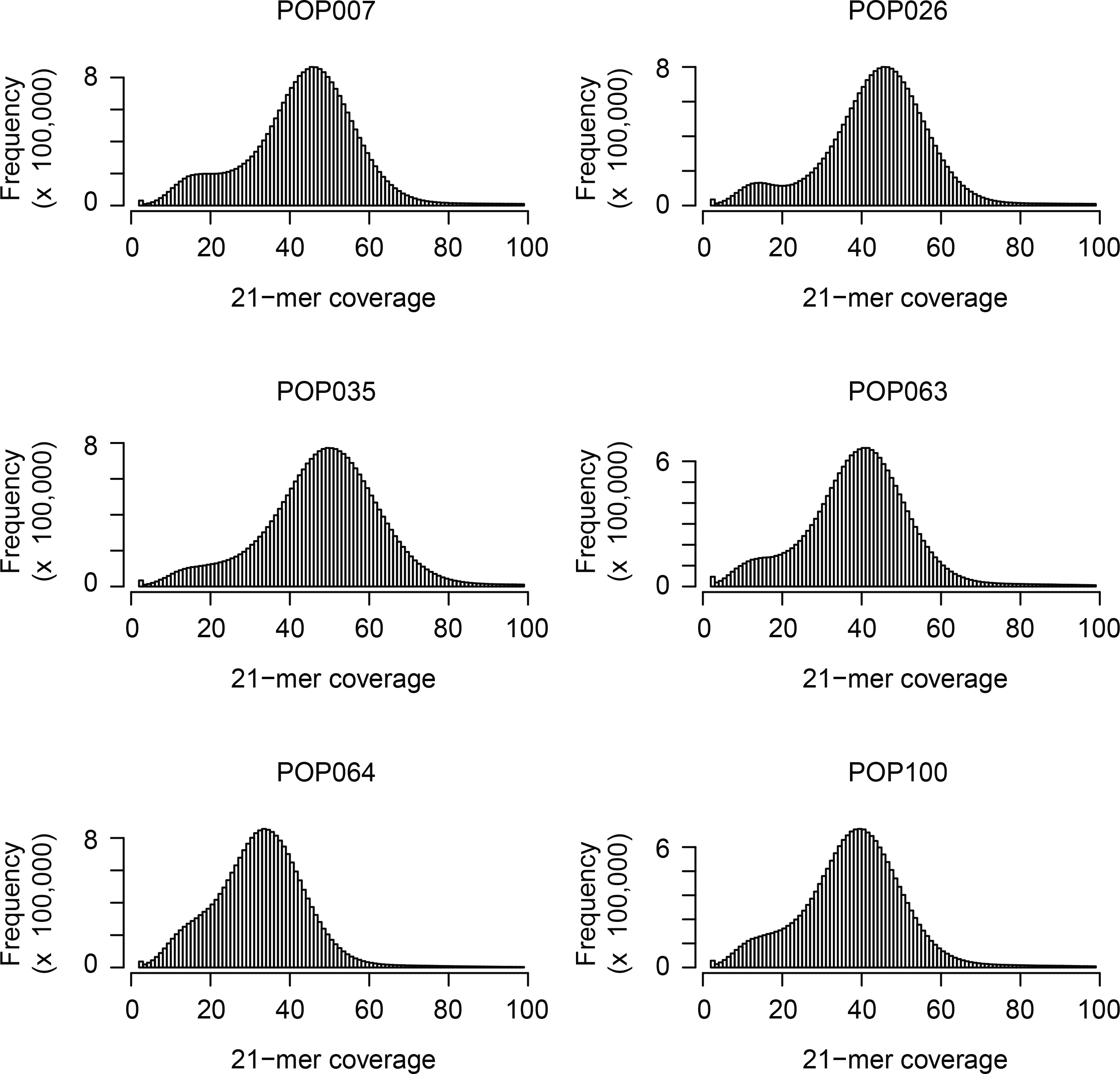
**Distribution of unique 21-mers in whole-genome resequencing data.** The coverage of unique 21 nt k-mers is plotted for each of the six populations that underwent whole-genome resequencing. The first peak in coverage represents 21-mers found in only one of the two grandparents, while the second, more prominent peak represents those found in both.

**Figure S4.**
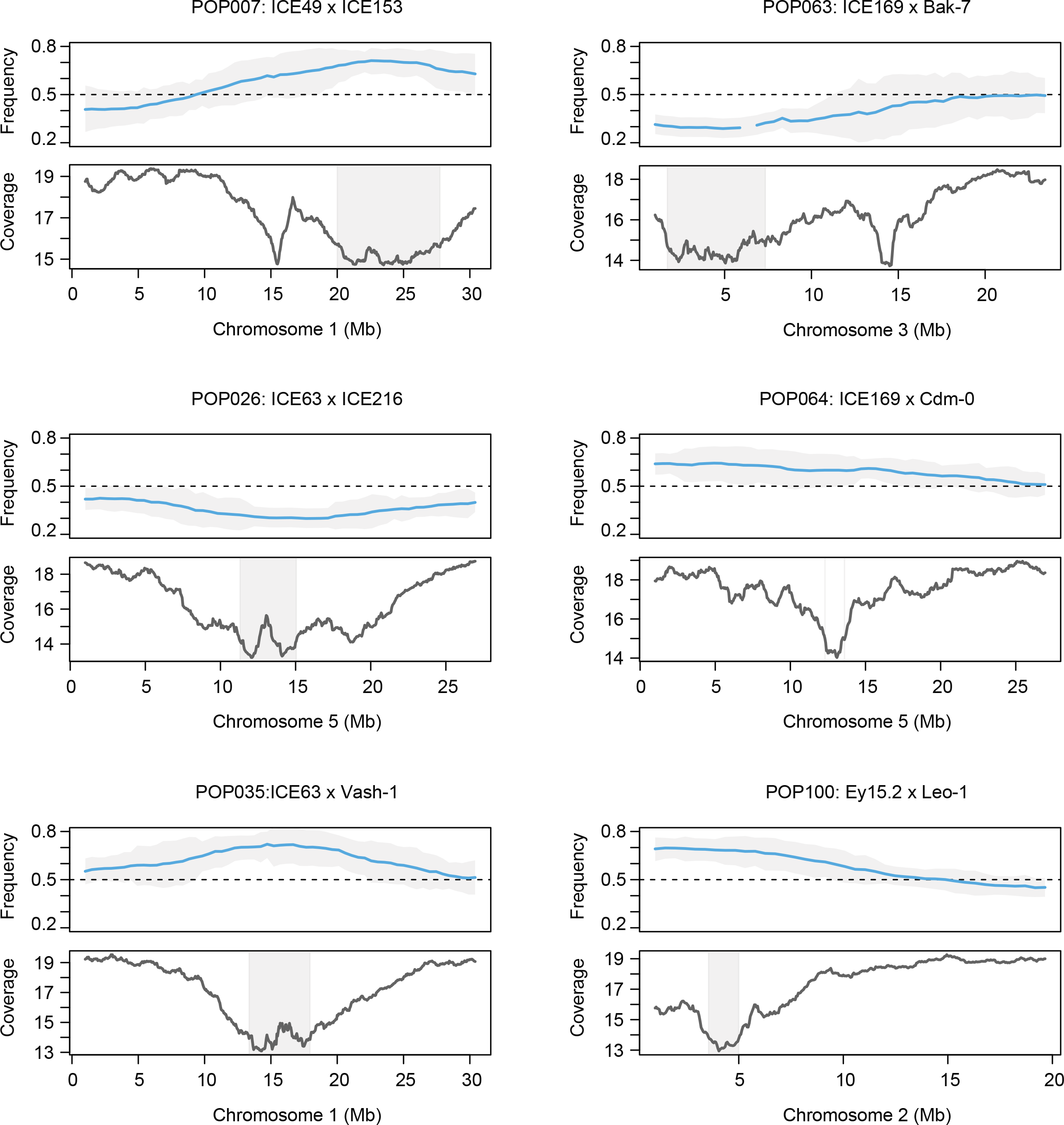
**21-mer coverage from whole-genome resequencing can be used to refine mapping intervals.** For each population, the upper panel displays the beta-binomial modeled allele frequency estimates (blue) and their 95% confidence intervals (grey) as described in the legend for Figure 2. In the lower panel, the coverage of 21-mers unique to only one of the two grandparents (coverage < 25x) is plotted in 1 Mb sliding windows (50 kb steps). Coverage decreases in the candidate regions. Intervals (grey box) are defined by merging windows with values within 1x coverage of the minimal window in each population. No candidate region was defined for POP064 as coverage decrease coincides with the centromere, not the distorted region.

**Figure S5.**
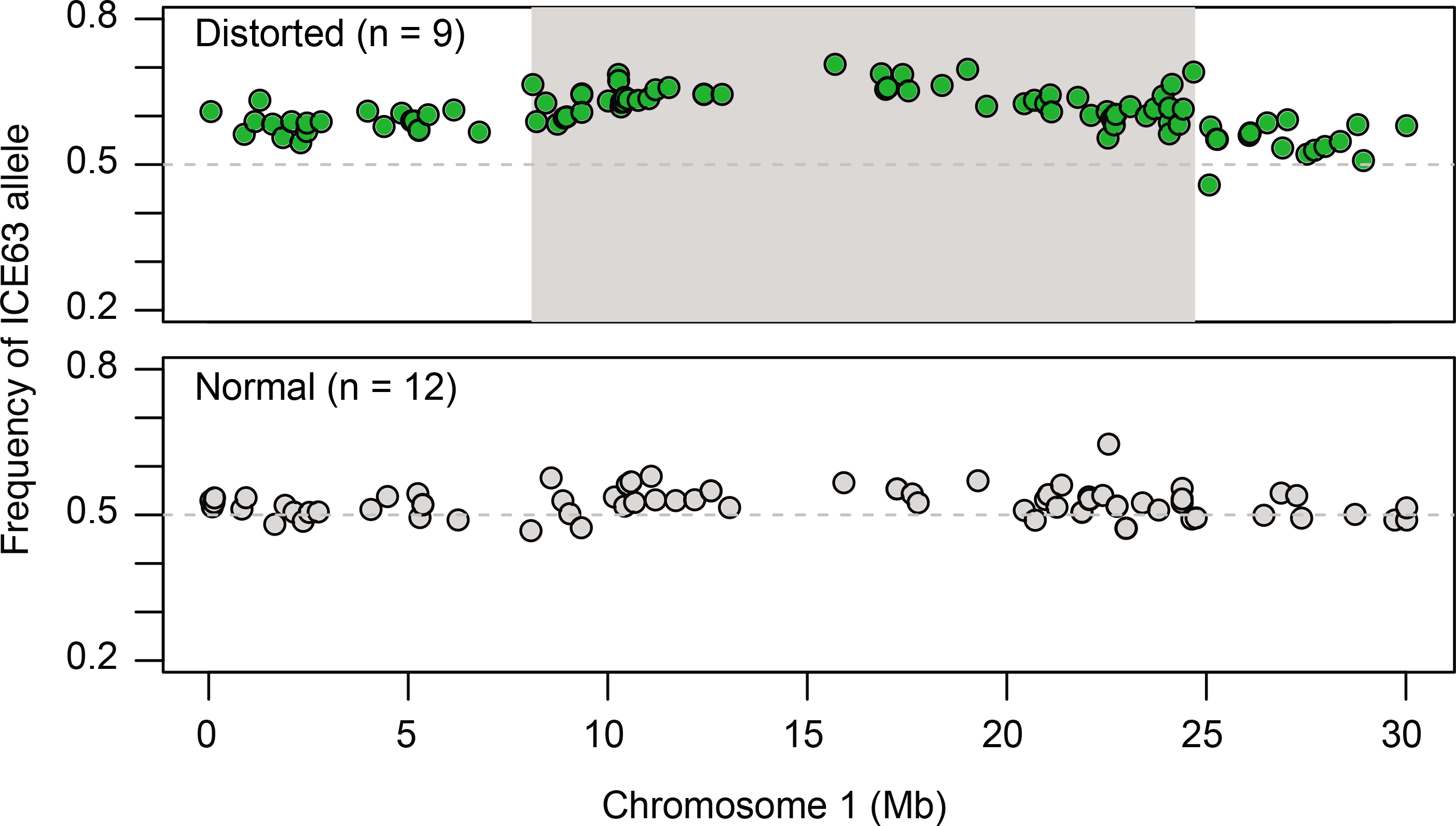
**Increasing the number of analyzed segregants can be used to refine mapping intervals.** Bulked segregant analysis was performed for grandparental accessions that repeatedly contributed distorted loci (Star-8 [Figure 6C], ICE63 [shown here], and ICE49). Sequencing reads were combined for populations exhibiting distortion or not exhibiting distortion when crossed to the focal grandparent. An average of over 800x coverage was achieved at sites segregating between the focal accessions and all other members in the bulk. A candidate interval (grey box) was obtained by merging all segregating positions within 5% of the maximal allele frequency. Data for ICE49 not shown, as there were too few segregating sites.

